# Environmental filtering shapes divergent bacterial strategies and genomic traits across soil niches

**DOI:** 10.64898/2026.01.16.699881

**Authors:** Tim Goodall, Robert I. Griffiths, Bridget Emmett, Briony Jones, Amy Thorpe, Daniel S. Read, Susheel Bhanu Busi

## Abstract

Soil pH is a predominant factor in structuring microbial communities; however, its role in shaping microbial life-history traits across large spatial scales remains underexplored. Here, we hypothesised that bacterial ubiquity, or niche breadth, across a diverse collection of soils is linked to genomic traits. We leveraged a national-scale survey of UK soils (the Countryside Survey) and 16S rRNA gene sequencing data with trait annotations (estimated genome size, coding density, and rRNA operon copy number) to examine trait-environment-niche breadth relationships. Our analyses revealed that soil pH was the dominant environmental driver of niche classification and bacterial community traits along the niche range. Low pH soils (pH <5.5) hosted ubiquitous taxa with larger genome sizes, lower coding densities and lower rRNA copy numbers, implying slower growing taxa with higher genetic facilities. Mildly acidic soils (pH 5.5 to 7) favour higher rRNA copy numbers, intermediate genome sizes and moderate coding densities. Alkaline soils (pH >7) feature communities with the smallest niche range, smallest genomes and highest coding densities. Here, specialisation occurs through streamlining with simpler, smaller genomes favoured. We found that generalist taxa were widespread across the pH range, becoming dominant under acidic conditions, while taxa adapted to higher pH were comparatively scarce in their distribution. These findings identify soil pH as a key physiological filter that aligns microbial genomic traits and ecological strategies across landscapes. By extending prior site-specific results to a broad-scale context, our study highlights how trait-based metrics can predict microbial responses to soil conditions, with implications for understanding ecosystem carbon cycling and informing land management practices aimed at sustaining soil health in the future.

## Introduction

Soil pH is often regarded as a *master variable* in soil ecology, exerting a strong influence on microbial community composition and function (Fierer and Jackson 2006, Rousk, Baath et al. 2010, Griffiths, Thomson et al. 2011, Neina 2019). In diverse soils, pH affects nutrient availability and physiological constraints, thereby determining which microorganisms can thrive (Neina 2019). Bacterial taxa vary in their pH tolerance, and this environmental filtering is thought to drive broad patterns of microbial diversity and biogeography. Understanding how pH and other soil properties filter microbial communities is crucial, as microbes mediate key ecosystem processes such as decomposition and nutrient cycling (Falkowski, Fenchel et al. 2008, Schimel and Schaeffer 2012, Fierer 2017).

Microorganisms differ widely in their ecological strategies, which can be characterised by their genomic traits. Notably, genome size and ribosomal RNA (rRNA) operon copy number are key indicators of life-history strategies (Madin, Nielsen et al. 2020). Bacteria with large genomes tend to be metabolically versatile, allowing them to adapt to fluctuating or resource-poor environments. Conversely, smaller genomes are often associated with specialists adapted to stable, resource-limited niches, favouring growth potential over versatility (Giovannoni, Cameron Thrash et al. 2014). Similarly, higher rRNA copy numbers are associated with greater growth potential (Klappenbach, Dunbar et al. 2000, Roller, Stoddard et al. 2016), enabling rapid resource exploitation by taxa (Fierer, Bradford et al. 2007).

These trait patterns mirror classical ecological trade-offs: generalists (“opportunists” or r-strategists) invest in broad functional capacity and fast response, whereas specialists (K-strategists) may streamline their genomes for efficient resource use (Malik, Martiny et al. 2020). Importantly, such genomic traits have been linked to the range of conditions that a microbe can tolerate, that is, its ecological niche (Barberán, Ramirez et al. 2014). This can be quantified using Levins’ niche breadth index, which characterises how widely and evenly a taxon occupies environmental space (Levins 1968). Unlike simple occurrence frequency metrics, Levins’ index integrates both prevalence and proportional abundance across conditions, providing a more mechanistic estimate of ecological tolerance. High Levins’ values therefore denote broadly distributed generalists, while low values identify specialists confined to narrow environmental ranges. This quantitative framework offers a robust means of distinguishing microbial life history strategies, including niche-restricted specialists.

Previous studies have suggested that soil pH strongly influences microbial traits and distribution. For instance, in highly managed agricultural soils, pH has been identified as the primary driver of bacterial genomic traits and niche breadth (Goodall, Busi et al. 2025). Communities in lower-pH soils (pH 6 to 7) were favoured by bacteria with larger genomes, higher rRNA copy numbers, and greater ubiquity (broader niches). In contrast, higher-pH soils (above pH 7.5) hosted more specialised microbes with smaller genomes. This finding implies that tolerance to acidity may confer a broader distribution, potentially because acid-tolerant generalists can also persist in neutral soils, whereas alkaliphiles (high pH specialists) are confined to rarer habitats (Goodall, Busi et al. 2025). However, it remains unclear whether these pH–trait relationships hold true across much larger spatial scales with more varied land uses beyond agricultural settings, with their comparatively narrow pH range. Soil properties such as organic carbon and moisture often covary with pH in natural environments, which could modulate or confound the influence of pH (Griffiths, Thomson et al. 2011). A broader-scale assessment is needed to confirm that pH acts as a consistent ecological filter for microbial traits and to explore how other factors contribute to pH. Soils offer an ideal test case owing to their strong environmental gradients and well-characterised properties. National-scale monitoring (the UK Countryside Survey https://www.ceh.ac.uk/our-science/projects/countryside-survey) has demonstrated that soil habitats exhibit distinct, predictable ranges of pH, bulk density, and organic matter that reflect differences in land use, parent material, and climate (Emmett, Reynolds et al. 2010).

Here, we address this knowledge gap by examining microbial trait–environment relationships at the landscape scale. We use data from the UK Countryside Survey, which encompasses hundreds of soils across different habitats and management intensities, providing a wide gradient in pH and other soil properties. For each soil bacterial community, we derived trait measures including the community-weighted mean genome size, rRNA operon number, coding density, and the Levins’ niche breadth of the constituent taxa. We hypothesised that bacterial ubiquity across sites is linked to genomic traits conferring adaptation to soil pH conditions. By extending prior findings on pH as a master variable, our study integrates genomic trait diversity and niche breadth to elucidate how environmental filtering shapes microbial life-history strategies at large scales. This trait-based perspective provides insight into the ecological assembly of soil microbiomes and their potential functional implications for ecosystems.

## Methods

### Sample collection and physicochemical measurements

Soils were collected as part of the 2007 UK Countryside Survey (https://www.ceh.ac.uk/our-science/projects/countryside-survey) following published guidelines (Maskell 2008). The Countryside Survey is a long-running survey of the UK landscape and is designed to encompass multiple sites from each of the recognised land classes of the UK. Briefly, a clean, unused plastic tube of 5 cm diameter was used to collect soil core samples from the top 15 cm at each sample site; cores were sealed into pre-labelled plastic bags to prevent the transfer of soil residue between samples. The cores were transferred to the laboratory on the day of collection and subjected to multiple key analyses (BA Emmett 2008). These included measurements for organic matter through loss on ignition, percent carbon, percent nitrogen, moisture, and pH.

For molecular analyses, cores were frozen at −20°C until processing. The cores were lightly defrosted, and a sub-sample of soil collected from below the organic horizon (thus excluding fine roots) was homogenised and archived at −20°C prior to DNA extraction.

### DNA extraction, PCR and sequencing

DNA was extracted from 0.25 g of the frozen, archived soil. Briefly, soil was weighed by means of pre-sterilised (immersion in 5% bleach and 70% ethanol wash) apparatus into Powersoil DNA 384 Isolation Kit (Qiagen Ltd.) plates, and the DNA was extracted according to the manufacturer’s instructions. Samples were randomly distributed across extraction plates, and each extraction plate incorporated negative extraction controls. PCR and sequencing were performed as per Jones et al. (Jones, Goodall et al. 2021), with the samples split equally across three Illumina MiSeq V3 600 cycle flow cells to maintain a sequencing depth target of >10k reads per sample.

### Bioinformatic analysis and ASV filtering

The forward reads of each of the three demultiplexed sequencer outputs where independently processe dusing DADA2 (Callahan, McMurdie et al. 2016). Filtering settings were truncLen=280, maxN=0, maxEE=1. Amplicon sequence variants (ASVs) were inferred, chimeras removed, and sequence tables constructed. These were then merged by ASV sequence and the assign Taxonomy function with bootstrapping set to 80 was used to assign each ASV it’s SILVA 138.2 identity (Quast, Pruesse et al. 2013).

Samples with <5,000 reads were removed from the dataset, and ASVs with taxonomic identities of chloroplast/mitochondrial assignments, or non-bacterial domains were removed. Remaining samples were rarefied to 4,369 reads, with a total of 26,372 filtered bacterial ASVs recorded across 899 samples. To permit the calculation of niche breadth, the rare taxa, with relative abundance <0.01% and occupancy <10%, were removed. Removal of these rare taxa produced a shortlist of 1,036 taxa; these filtered datasets were used in all subsequent analysis.

### Genomic trait assessment and Levins’ niche breadth

Genome size, coding density and rRNA operon copy number were estimated by linking each of the filtered 1,036 ASVs taxonomic assignments to the closest available representative genome in the GTDB database (Parks, Chaumeil et al. 2025). Where species-level matches were unavailable, values were assigned hierarchically at the genus, family, or higher taxonomic ranks following established trait-propagation approaches (Langille, Zaneveld et al. 2013). The largest proportion of matches, 65%, were to Family level or above, 18% to Order, 6% to Class and 9% to Phylum (Supp. Fig. 1). Where multiple database entries per ASV were found the median vales were calculated and used in subsequent analysis. rRNA operon copy number accuracy was refined using the machine-learning predictor trained on complete genomes, ANNA16 (Miao, Chen et al. 2024), improving estimates for taxa lacking curated entries. The environmental space was defined by calculating the hypervolume occupied by the individual taxa. To determine this, the *hypervolume* R package was used (Blonder, Morrow et al. 2018) which factored in the following metadata variables: pH, percent carbon, percent nitrogen, loss on ignition (LOI), C:N ratio, and moisture content. Subsequently, the estimated hypervolume space was divided into 10 bins based on k-means clustering. These bins were used to calculate per ASV niche breadth scores (Levins’ index) from relative abundance distributions across all samples using the R package *MicroNiche* (Finn, Yu et al. 2020). The Levins’ index provides a measure of how evenly a taxon occupies the hypervolume-environmental space, with higher values indicating broader environmental tolerance and generalist behaviour, and lower values reflecting restricted specialists.

### Statistical modelling and data analysis

Each sample’s weighted means were derived by computing per-ASV abundance-weighted community means (CWMs) for genome size, coding density and rRNA copy number, and Levins’ niche breadth score. These weighted means were also calculated for each taxonomic phylum (PWMs) across the samples, enabling the examination of broad taxonomic trends.

We employed statistical models to test the influence of traits and environment on niche breadth. Generalised Additive Models (GAMs, using the *mgcv* package in R (Wood 2011)) were used to allow for non-linear relationships between predictors (traits or environmental variables) and the niche breadth response. For each model, we assessed the proportion of variance (deviance) explained to evaluate model fit. We also performed variance partitioning using LMG *relative importance* analysis (with the *relaimpo* package (Groemping 2006)) to determine the contribution of each predictor to explained variance. To ensure robustness of our findings, we conducted permutation tests (n = 999 permutations) for key model results, assessing the significance of trait effects against a randomised null distribution.

We constructed and evaluated models at two different organisational scales: (i) the individual taxon (ASV) scale, examining whether an ASV’s genomic traits predict its own niche breadth; (ii) the community trait scale, relating community-weighted mean trait values to the overall niche breadth of the community. Model comparisons across these scales allowed us to discern the relative importance of intrinsic genomic traits in shaping niche breadth. pH change-points for phyla present in at least 90 samples (∼10%) were assessed using the R package *TITAN2* (https://github.com/dkahle/TITAN2). Figures were generated in R using ggplot2 (https://ggplot2.tidyverse.org).

### Code and data availably

Countryside survey 16S sequences are deposited in the European Nucleotide Archive, accession PRJEB45286 and wider metadata for the samples is available at https://catalogue.ceh.ac.uk/documents/79669141-cde5-49f0-b24d-f3c6a1a52db8.

R-scripts used in the analysis of this data is available from ZENODO, DOI#############

## Results

### pH as the dominant determinant of soil-environment niches

To quantify environmental niche breadth, we constructed a multivariate environmental hypervolume using the edaphic variables pH, %C, %N, C:N ratio, LOI and %moisture (*see Methods*). This hypervolume represented the realised environmental space sampled across all soils. Prior to estimating Levins’ scores, we assessed the contribution of the edaphic variables to hypervolume space using a Random Forest classification of niche bin identity. Soil pH emerged as the most important predictor of environmental bin assignment, far exceeding the influence of C:N ratio, moisture, LOI, or nutrient concentrations (Fig. 1). This confirmed pH is the dominant axis along which environmental space is partitioned, reinforcing its role as a primary environmental filter structuring microbial distributions.

**Figure 1.**
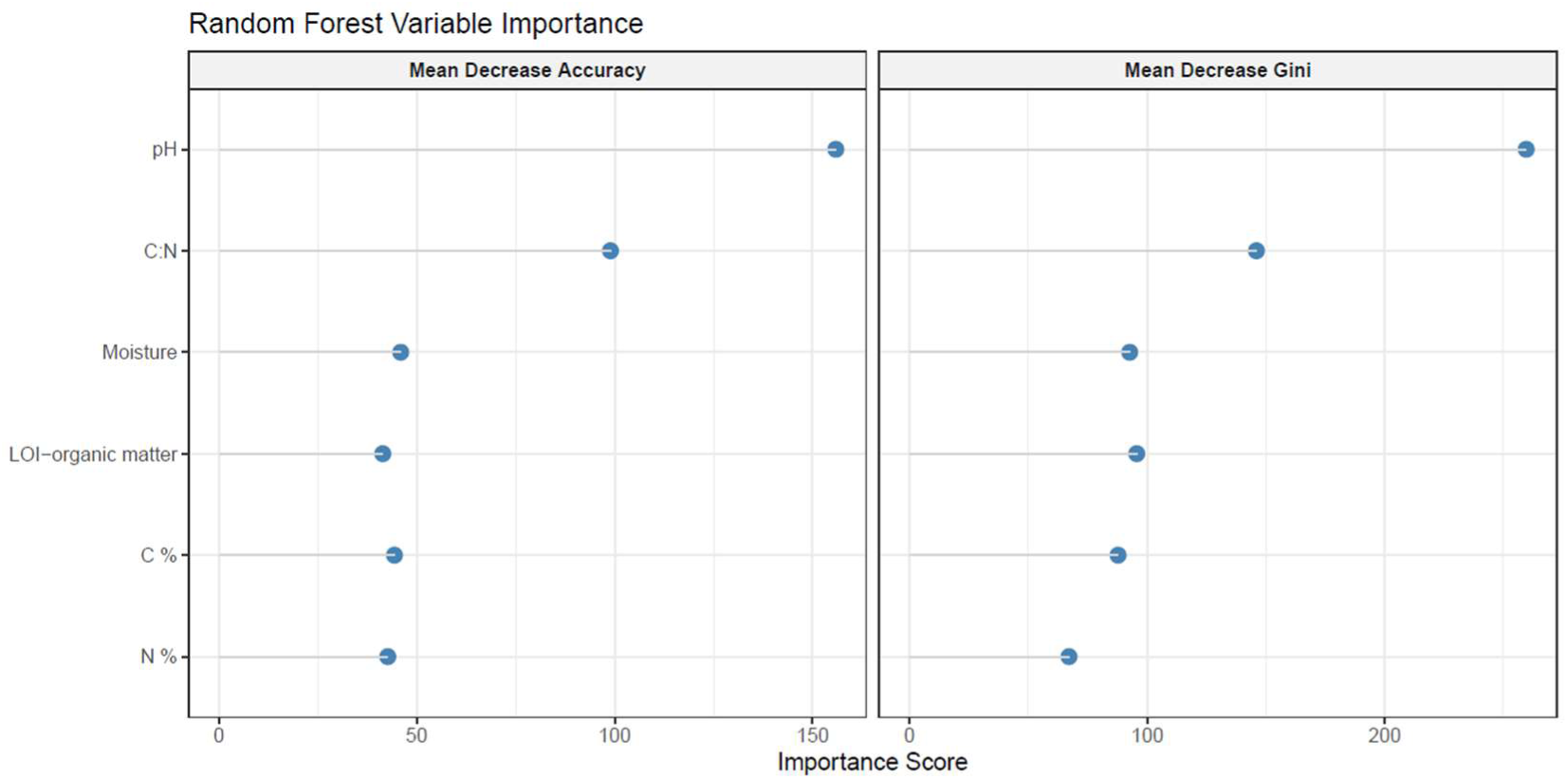
Random Forest ranked importance of soil environment variables in niche bin classification.

#### Minimal influence of genomic traits on ASV niche breadth

To test whether genomic traits act as intrinsic determinants of ecological strategy, we first evaluated whether they could predict microbial niche breadth at the level of individual taxa. If niche breadth is a direct property of a taxon’s genomic traits, environmental distributions should be apparent. At the ASV level, genomic traits showed minimal explanatory power for ecological niche breadth. The GAM relating Levins’ niche breadth to genome size, coding density, and rRNA operon copy number explained <5% of model deviance, with a weaker fit of 4% variance observed in LMG partitioning analyses (Supp. Tab. 1). Genome size and rRNA copy number showed no detectable association with niche breadth (Fig. 2a-b). In contrast, coding density displayed a weak but statistically significant negative relationship with Levins’ index (Fig. 2c), indicating that taxa with broader environmental distributions tend to have slightly lower coding densities. Although the effect size was small (R² = 0.03), this pattern is consistent with the idea that generalist taxa may maintain larger proportions of non-coding or regulatory DNA, potentially supporting greater ecological flexibility.

**Figure 2.**
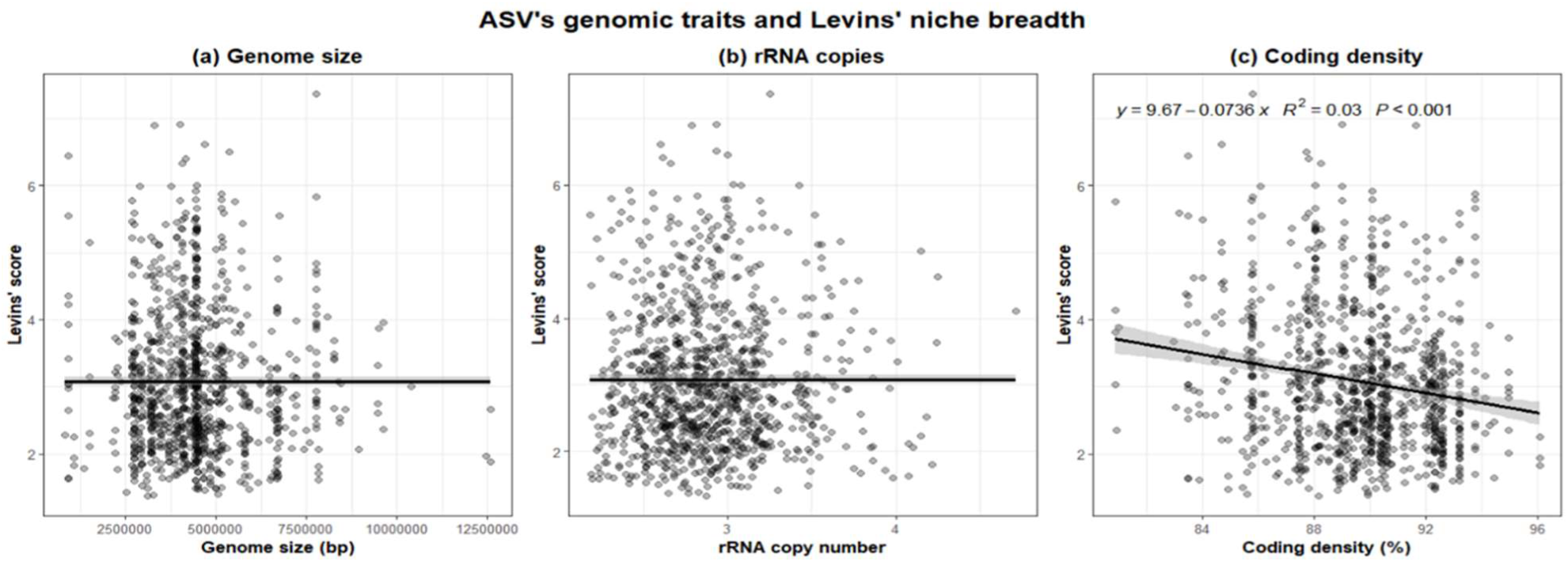
Relationship between each ASV’s Levins’ niche breadth score and genomic traits (a) Genome size, (b) rRNA copies, (c) Coding density with regression values.

Overall, these results suggest that while coding density shows a modest association with realised niche breadth, core genomic traits alone cannot reliably distinguish narrow-ranged specialists from broadly distributed generalists in our dataset. The absence of strong or consistent relationships indicates that broad genomic functional potentials do not, on their own, determine a taxon’s realised environmental distribution. Instead, niche breadth at the ASV level is likely shaped by ecological history, environmental filtering, and underlying phylogenetic constraints rather than any single genomic feature.

### Trait–niche structure at the community level

The absence of predictive power at the ASV level suggested that genomic traits do not determine niche breadth in isolation. Therefore, in line with emergent properties principles, we hypothesised that ecological filtering may lead to trait distributions across taxa detectable at the scale of communities (Fierer, Barberán et al. 2014). When genomic traits were aggregated into community-weighted means (CWMs), clear and significant associations between trait composition and realised niche breadth emerged that were not evident at the ASV scale (Fig. 3). Community-weighted coding density emerges as the dominant predictor of Levins’ niche breadth, followed by rRNA copies and genome size. The combined genomic traits explained 70% of deviance in GAMs and 62% of variance in LMG partitioning analyses (Supp. Tab. 2).

**Figure 3.**
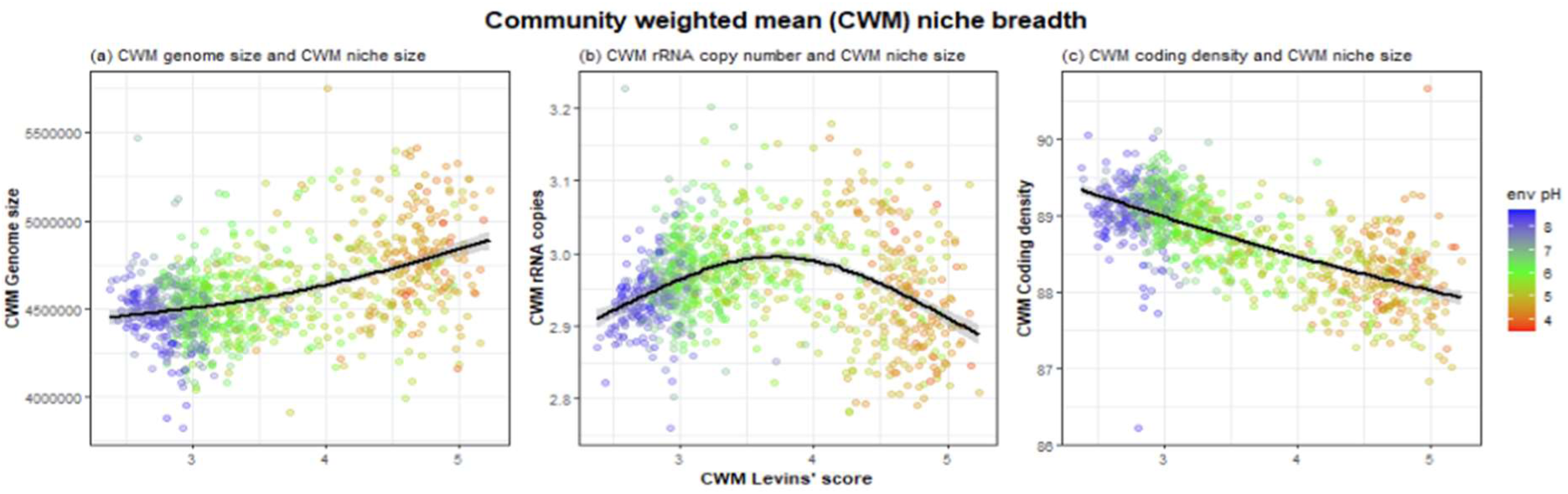
Community weighted mean relationships between niche breadth and genomic traits, (a) Genome size, (b) rRNA copies, (c) Coding density. Points coloured by sample’s pH.

Using CWM values we reveal a breakdown, at the community level, in the canonical proviso that larger genomes are linked to higher rRNA copy number when soils are < pH ∼5.5 (Fig. 3, orange and red point colour). These soils harbour communities with CWMs featuring larger genomes with lower coding density, yet they also have low rRNA copy numbers conferring the slowest growth potential and adaptation through genomic regulatory complexity and size. In the middle ground, pH ∼5.5-7.5 (Fig. 3, green point colour), CWMs show mid genome size / coding density with high rRNA copy numbers, suggesting growth rate as a driver for adaptation. Whilst under basic conditions, > pH ∼7.5 (Fig. 3, blue point colour) CWMs feature smaller genomes, high coding density and low rRNA copy number, suggesting that under these edaphic conditions genomic streamlining is prevalent. Communities from acidic soils also show the highest Levins’ niche breadth scores implying taxa dominant at pH<5.5 are more ubiquitous across niche space but proliferate at low pH at the expense of range limited taxa that favour pH above ∼5.5.

### CWM genomic traits across the broader environmental habitat

Broad habitat classifications from the Countryside Survey (Fig. 4) further demonstrate that ecosystem context structures the realised CWM niche breadth and genomic traits across soils. Habitats classifications that strongly feature samples from the pH zone ∼5.5-7.5 (Fig. 4, point colour green), Improved and Neutral Grasslands, Arable and Horticulture and Urban, exhibited lower CWM Levins’ niche breadth values, smaller CWM genome size, and higher CWM coding density and rRNA copies. These patterns suggest dominance by more ubiquitous, fast growing copiotrophs (r-strategists), aligning with elevated nutrient turnover, and recurrent disturbance through anthropogenic manipulation. In contrast, in the acidic habitats, such as Bog and Coniferous Woodland where pH is rarely above 5.5 (Fig. 4, point colours orange to red), CWMs show reduced niche breadth, larger genome size and coding density, yet lower rRNA operon copies. This reflects strong environmental filtering and the dominance of specialist-adapted, ubiquitous, oligotrophic taxa (k-strategists).

**Figure 4.**
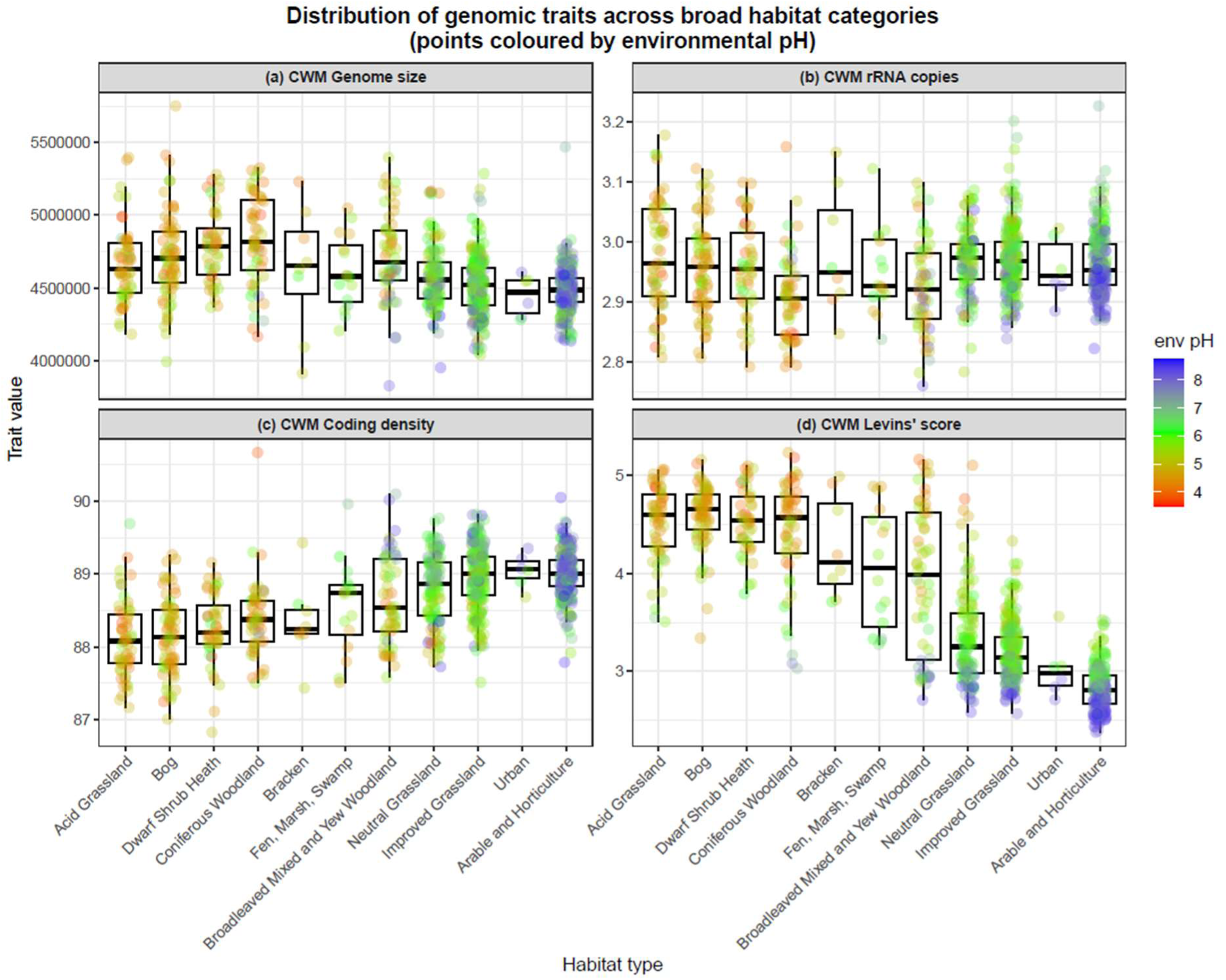
Box plots of samples assigned to their broad habitat classification for community weighted mean (CWM) genomic traits – (a) CWM-Genome size (bp), (b) CWM-rRNA copies, (c) CWM-Coding density (%), and (d) CWM-Levins’ niche breadth score. Points coloured by sampled community’s soil pH.

### Community compositional effects

Community composition along the pH gradient is visualised at the phylum level in Figure 5. Below pH 5.5, the abundance of members of Actinomycetota, Bacillota, and Bacteroidota decline with decreasing pH, whereas the ubiquitous phylum Acidobacteriota increases in abundance. To assess how phylogenetic organisation may inform genomic traits, we estimated the phylum-weighted mean (PWM) values across all samples. Our analysis demonstrates that phylogenetic trait strategies differ markedly across dominant phyla (Fig. 6). For example, the contrasting PWM-coding density (higher in Acidobacteriota versus lower in Bacillota), and PWM-rRNA copies (increased in Bacillota versus decreased in Acidobacteriota), suggesting maximum growth potential as an important part of this phylum’s survival and proliferation strategy.

**Figure 5.**
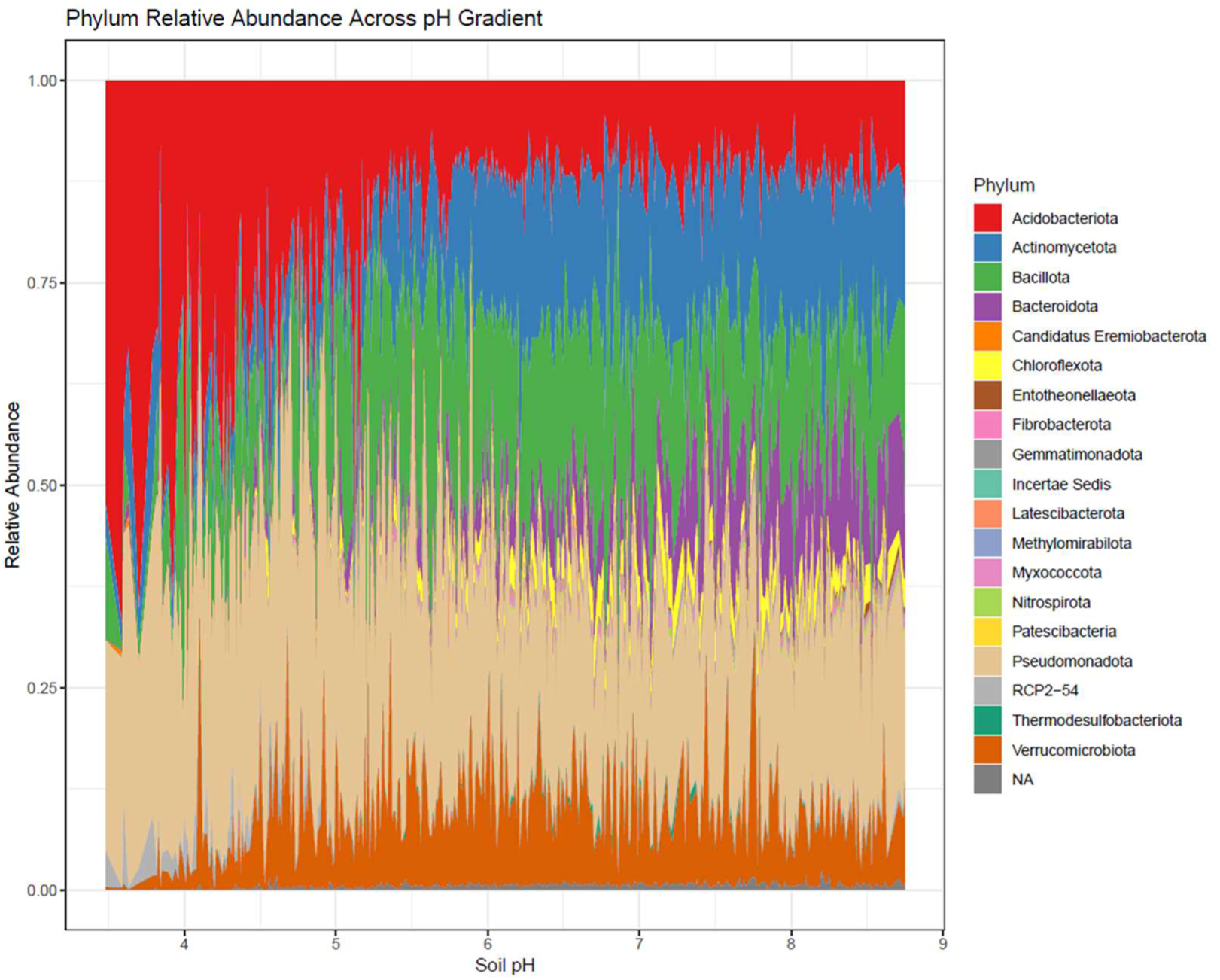
Plot of abundance of bacterial phyla along the sampled pH gradient.

**Figure 6.**
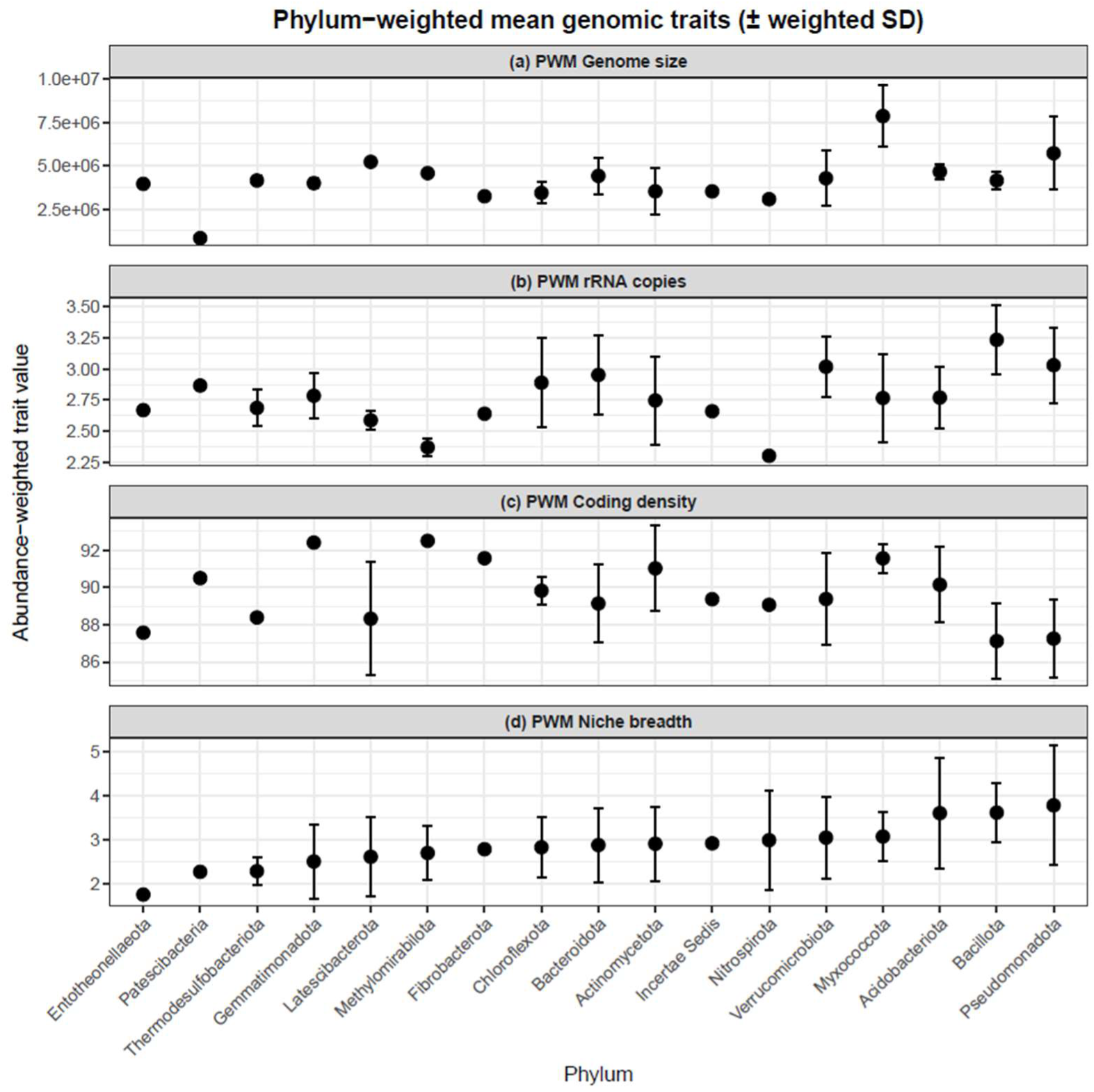
Phylum weighted mean (PWM) genomic traits of bacterial phyla, (a) PWM-Genome size (bp), (b) PWM-rRNA copy number, (c) PWM-Coding density (%), (d) PWM-Levins’ niche breadth. Point height represents the weighted mean; top and bottom whiskers ± abundance-weighted standard deviation across ASVs within each phylum.

To understand how the master variable (pH) influences these strategies, the relative abundance pH change points for phyla were assessed (*Methods*). We found that at the phylum level, there is an important community change between pH 5 and 5.4 (Fig. 7, blue dashed lines where decreasing taxa respond most strongly, red dashed line where increasing taxa respond most strongly). The dominant taxa of Actinomycetota, Bacteroidota, Chloroflexota, and Verrucomicrobiota showed marked increases in their abundance before or straddling these community change points, while importantly the ubiquitous phyla of Acidobacteriota decreased. These pH driven community compositional shifts indicate that members of these large and diverse phyla are key contributors to the observed CWM value changes.

**Figure 7.**
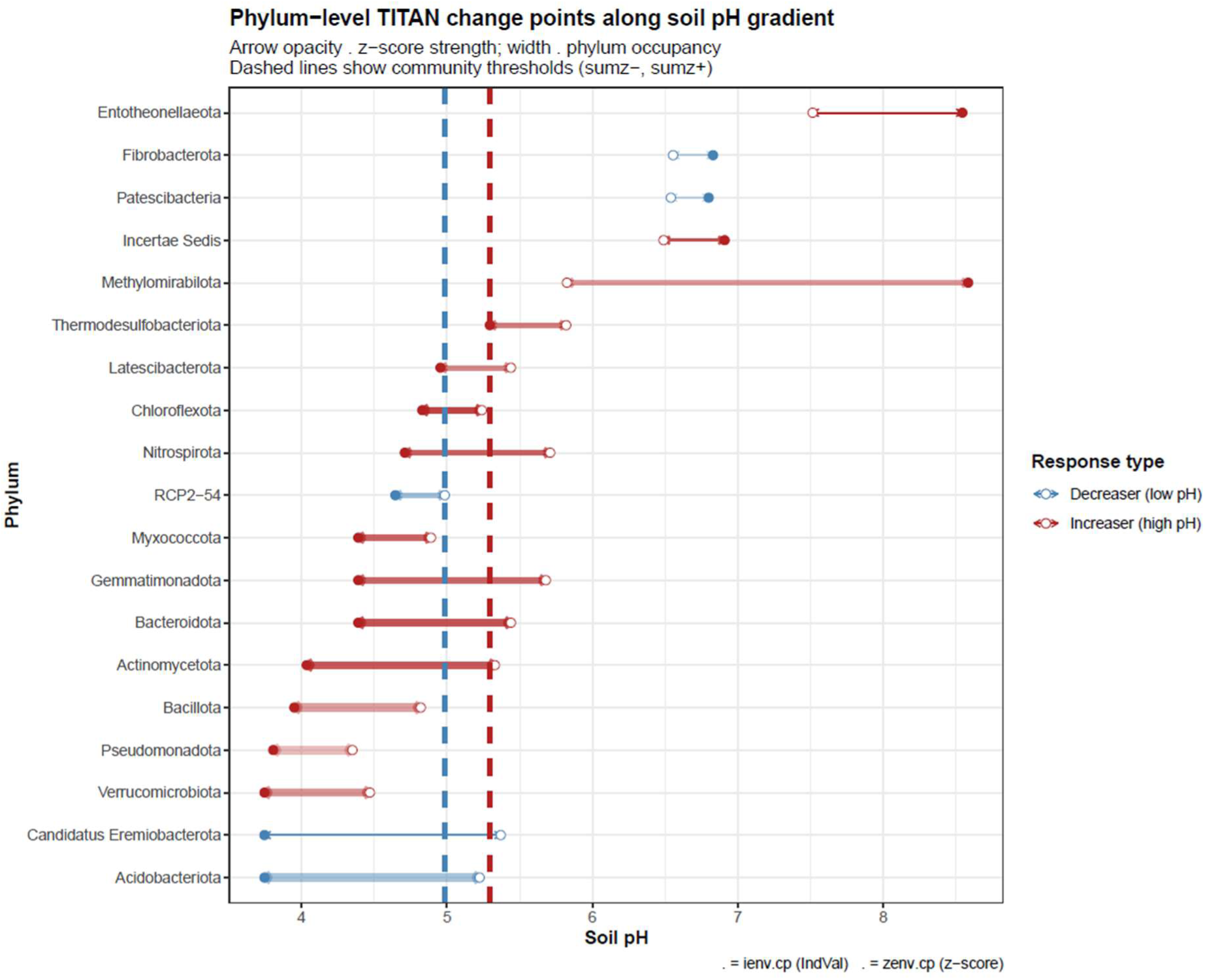
Threshold indicator taxa analysis (TITAN) of Phylum level abundance change-points along the pH gradient. Arrow opacity illustrates z-score strength, arrow width represents phylum occupancy, and colour indicates whether phyla increase (red) or decrease (blue) along the gradient. Dashed lines show the community change points for increasers (red dashed) and decreasers (blue dashed).

## Discussion

Our findings establish soil pH as a key environmental filter that aligns microbial traits and distribution patterns at broad spatial scales. Even amid the complexity of diverse soil types and land uses, pH was the overriding factor structuring bacterial community niche space reinforcing its status as a master variable in soil microbial ecology. Notably, pH explained more variation in niche space than soil organic matter or nutrient ratios and levels, highlighting that pH-driven physiological constraints often supersede the direct effects of resource quantity and quality (Fierer and Jackson 2006, Rousk, Baath *et al*. 2010, Malik, Puissant et al. 2018). This result is consistent with emerging evidence that pH exerts a fundamental influence on microbial life in soil, directly affecting cell physiology, for example enzyme function (Sinsabaugh, Lauber et al. 2008, Puissant, Jones et al. 2019), membrane stability (Wei, Shan et al. 2023) and indirectly shaping nutrient availability (Weil and Brady 2017) and toxicity (Kinraide 1991). By acting as a strong environmental sieve, pH determines which bacteria can persist and how they strategise to survive.

Trait relationships become apparent at the community level (Piton, Legay et al. 2020) in-line with emergent properties principles (Konopka 2009, van den Berg, Machado et al. 2022) where community-level patterns arise from interactions among constituent members that cannot be deduced linearly from individual ASV propertied. Changes in the abundance of dominant taxa along the pH gradient drive these emergent traits (Supp. Fig. 2). For example, members of Acidobacteriota and Pseudomonadota, prevalent in soils below pH 5.5, display a decrease in their coding densities as they become more dominant.

Combined, these factors of dominance and adaptation help to explain why community weighted means reveal strong relationships. Coding density emerged as the most important CWM genomic trait related to niche breadth, both at the ASV level and more significantly at the community level. Our study revealed that communities featuring low-coding density, i.e. complex genomes incorporating regulatory complexity, have the largest niche breadth. Conversely, in high coding density communities, where genomes are more streamlined and lack regulatory complexity, niche range is more limited. This implies that as pH increases into high alkali condition, oligotrophy may be selecting for communities that maintain high coding density due to selection for growth economy (Giovannoni, Cameron Thrash et al. 2014). Our analysis shows that low-pH soils (< ∼5.5) harbour bacterial communities with traits indicative of niche generalisation, and that the exclusion of other taxa is bought about by a lack of genomic regulatory complexity. Communities in these acidic environments were enriched in taxa featuring larger genomes, low coding density and low rRNA operon counts (i.e. high genomic potential and complexity with low growth potential). These traits likely confer a competitive advantage in challenging conditions – an acid soil can impose multiple stresses (nutrient limitations, metal solubility issues, etc.), and a broader metabolic toolkit helps microbial communities utilise diverse resources and cope with variability (Wang, Yu et al. 2023). Larger genomes may carry genes for stress tolerance, versatile metabolism, and efficient resource acquisition.

Communities within mildly acid to neutral soils (pH ∼5.5–7) were composed of taxa with the highest rRNA operon counts, middling genome sizes along and coding density, suggesting dominant taxa here are “jacks-of-all-trades”, lacking extreme specialisation yet able to rapidly respond to opportunity. This observation aligns with previous reports that metabolic versatility (often correlated with genome size) is higher in communities under mildly-acidic pH conditions (Goodall, Busi et al. 2024).

In contrast, communities in neutral to alkaline soils (pH >7) exhibited traits of specialism through genomic streamlining, i.e. smaller genomes, highest coding density and fewer rRNA operons (Giovannoni, Cameron Thrash et al. 2014). Alkaline conditions, which can coincide with typically low levels of organic matter, yet with high pH mediated resource availability and resource quality (low C:N), appear to permit more streamlined bacterial genomes. It is possible that here, under favourable pH conditions, other edaphic factors such as nutrient resource availability increase in importance in shaping CWM genomic traits. For instance, Chuckran *et al*. (Chuckran, Hungate et al. 2021) reported that nutrient limitations often lead to variable traits, including genome size, GC content and 16S rRNA copies among bacteria.

This trait and distribution turnover parallels a functional threshold identified previously by Malik et al. (Malik, Puissant et al. 2018) who noted that above pH ∼6.2, soil communities undergo a significant change, with increased carbon use efficiency and dominance of organisms with smaller genomes. Our findings mirror this threshold-driven transition: as pH rises into the neutral range and beyond, bacterial communities shift toward a different regime of life-history traits. Together these findings indicate that soil properties mediate a shift from more adaptable generalist-dominated communities in acidic to mildly acidic habitats, where spatial heterogeneity is greater (Supp. Fig. 3), to more streamlined, specialised communities at high pH where resources have lower spatiovariation.

The ecological relevance of these patterns extends beyond pH to the structure of the habitats themselves. When communities were grouped by broad habitat classes (Bunce 1999), niche breadth and genomic traits varied, with highly managed neutral to basic habitats demonstrating specialisation through streamlining (lowest CWM niche breadth and very high coding density). While spatially complex, naturally acidic habitats (bogs, heathland etc.) demonstrate dominance by broad niche taxa with large genetic complexity and low growth potential.

Soil microbial communities drive decomposition, nutrient mineralisation (Schimel and Schaeffer 2012), and soil organic matter formation (Liang, Amelung et al. 2019, Buckeridge, Creamer et al. 2022) where shifts in pH and consequent microbial strategy could impact these processes through changes in substrate utilisation, growth rate and carbon use efficiency. From a land management perspective, our findings suggest that manipulation of soil pH could have predictable effects on the soil microbiome trait architecture. Agricultural practices like liming (to raise pH) or additions of organic amendments that acidify soil will shift the balance of microbial strategies. It is important to contextualise microbial trait distributions, not just taxonomic compositions, when evaluating a soil’s microbiome (Green, Bohannan et al. 2008, Malik, Martiny et al. 2020). In this way, genomic trait metrics may serve as sensitive indicators of soil microbial response trajectories. By linking environmental drivers to measurable traits and distribution patterns, we gain a mechanistic understanding of community assembly that complements traditional diversity surveys.

It should be born in mind that our findings are subject to certain limitations; primarily, while the GTDB database (Edwin, Fitzpatrick et al. 2024, Parks, Chaumeil et al. 2025) is comprehensive and actively curated, diverse environmental taxa may not have the best representation. Furthermore, in the absence of full-length 16S rRNA sequences, the hierarchical taxonomic assignment method used here inherently lacks fine-scale resolution. This underscores a critical need for alternative amplicon targets, such as *rpoB* (Ogier, Pagès et al. 2019), that facilitate higher-resolution amplicon sequence-based matching to genomic databases. The increasing availability of shotgun metagenomic data provides an essential next step to move beyond inferred traits, allowing detection of genes and pathways associated with pH tolerance, stress adaptation, nutrient cycling, membrane transporters, proton pumps, and carbon assimilation mechanisms (Bahram, Hildebrand et al. 2018). Integrating these functional profiles with trait-based frameworks can confirm whether the genomic signals we infer (e.g. genome size or rRNA copy number) reflect actual functional capacities. This validation is critical for building predictive models of how microbial traits translate into biogeochemical outcomes under changing soil conditions. Equally, incorporating the fungal component of soil communities, through metagenomics, will provide a more complete understanding of pH-driven microbial ecology as fungi can dominate decomposition and nutrient turnover in acidic or organic-rich soils (Strickland and Rousk 2010, Schneider, Keiblinger et al. 2012). Their enzymatic diversity allows the degradation of polymers like lignin and cellulose under conditions where bacterial activity is constrained by low pH (Rousk and Bååth 2011). Such multi-kingdom, metagenome-informed analyses could reveal whether pH exerts convergent evolutionary pressures across microbial domains or whether bacteria and fungi occupy complementary niches along the soil acidity spectrum.

Our results demonstrate that bacterial ubiquity in soils is linked to tolerance of pH variation, revealing environmental filtering as the dominant process structuring microbial communities and their genomic traits at broad spatial scales. Niche breadth emerges from the interaction between genomic trait composition and the strength of environmental constraints. Soil pH shapes not only which lineages persist, but also the balance between specialist and generalist strategies, and thus the functional potential embedded within communities. By identifying pH as a master filter that aligns genomic traits, niche breadth, and community-level strategy, this study provides a basis for predicting microbial responses to soil change, with implications for carbon cycling, nutrient turnover, and soil ecosystem resilience, because shifts in strategy from versatile generalists to specialists likely correspond to changes in substrate use, growth efficiency, and microbial carbon storage pathways.

---

**Supplementary figure 1.**
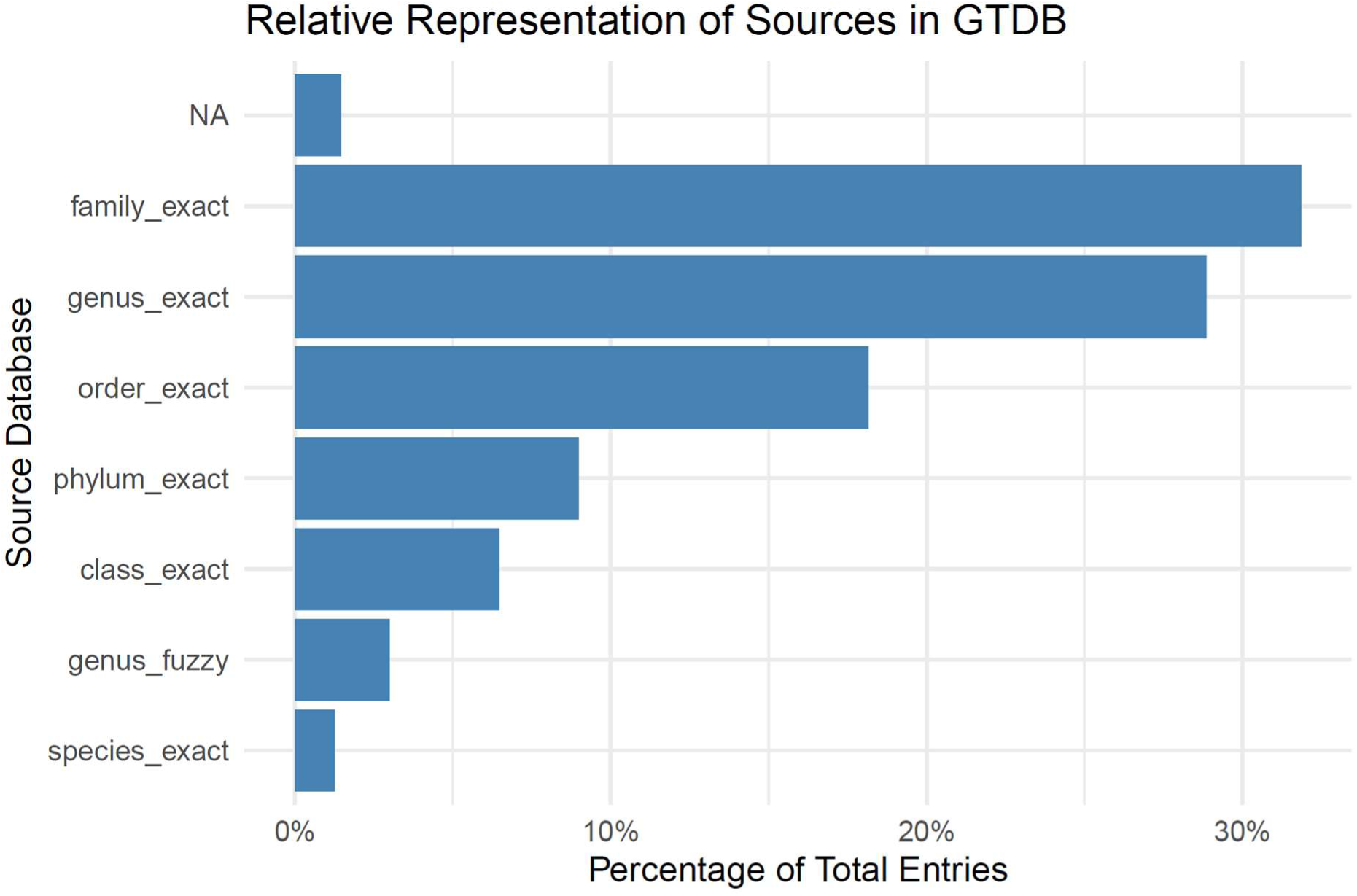
Taxonomic level of GTDB genomic trait assignments for filtered ASV.

**Supplementary figure 2.**
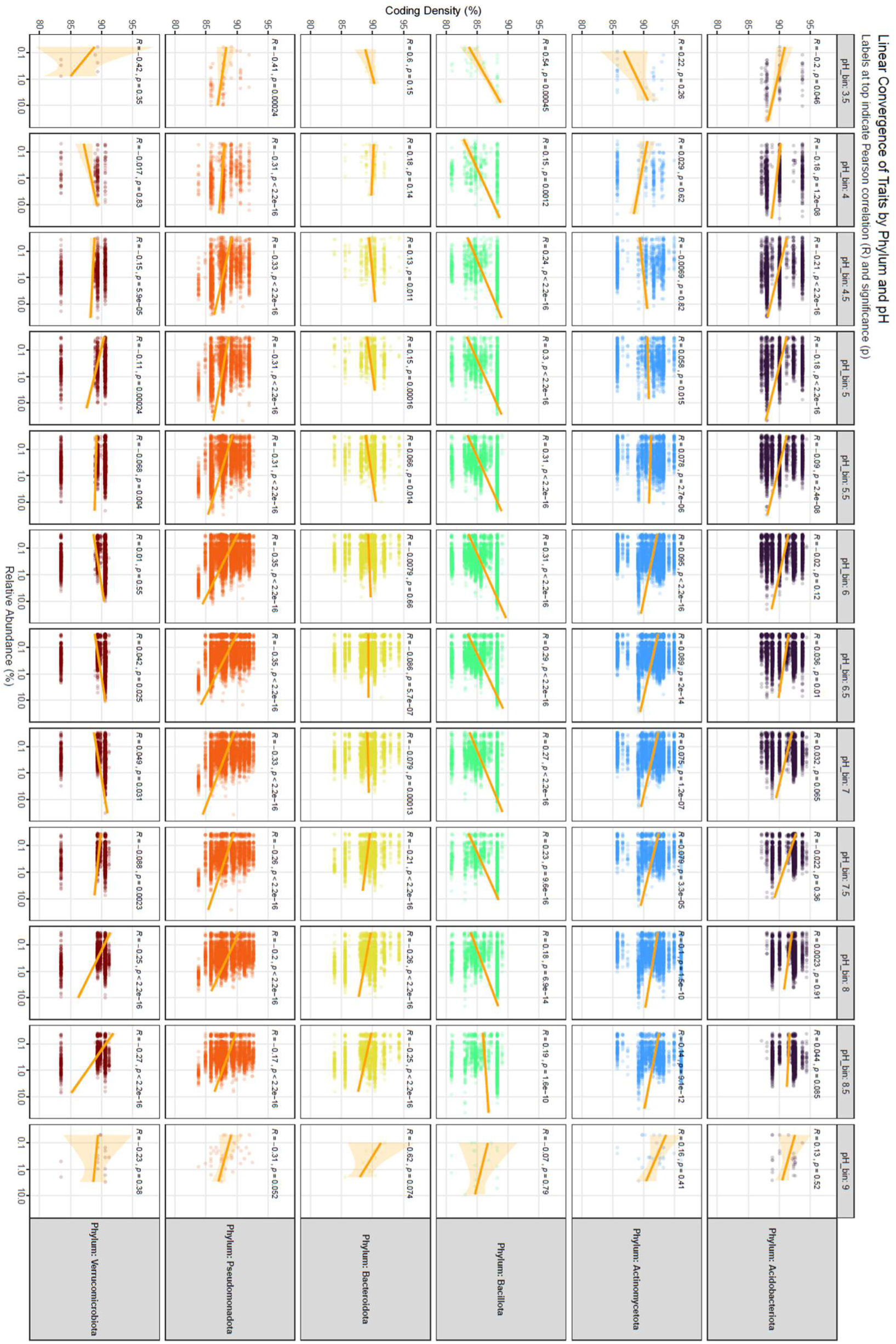
Taxon-specific trait convergence across the soil pH gradient. Relationship between ASV relative abundance and genomic coding density (CD) across major bacterial phyla and 0.5-unit pH intervals. Data are faceted by phylum (rows) and soil pH bins (columns) to illustrate within-group trait responses to environmental filtering

**Supplementary figure 3.**
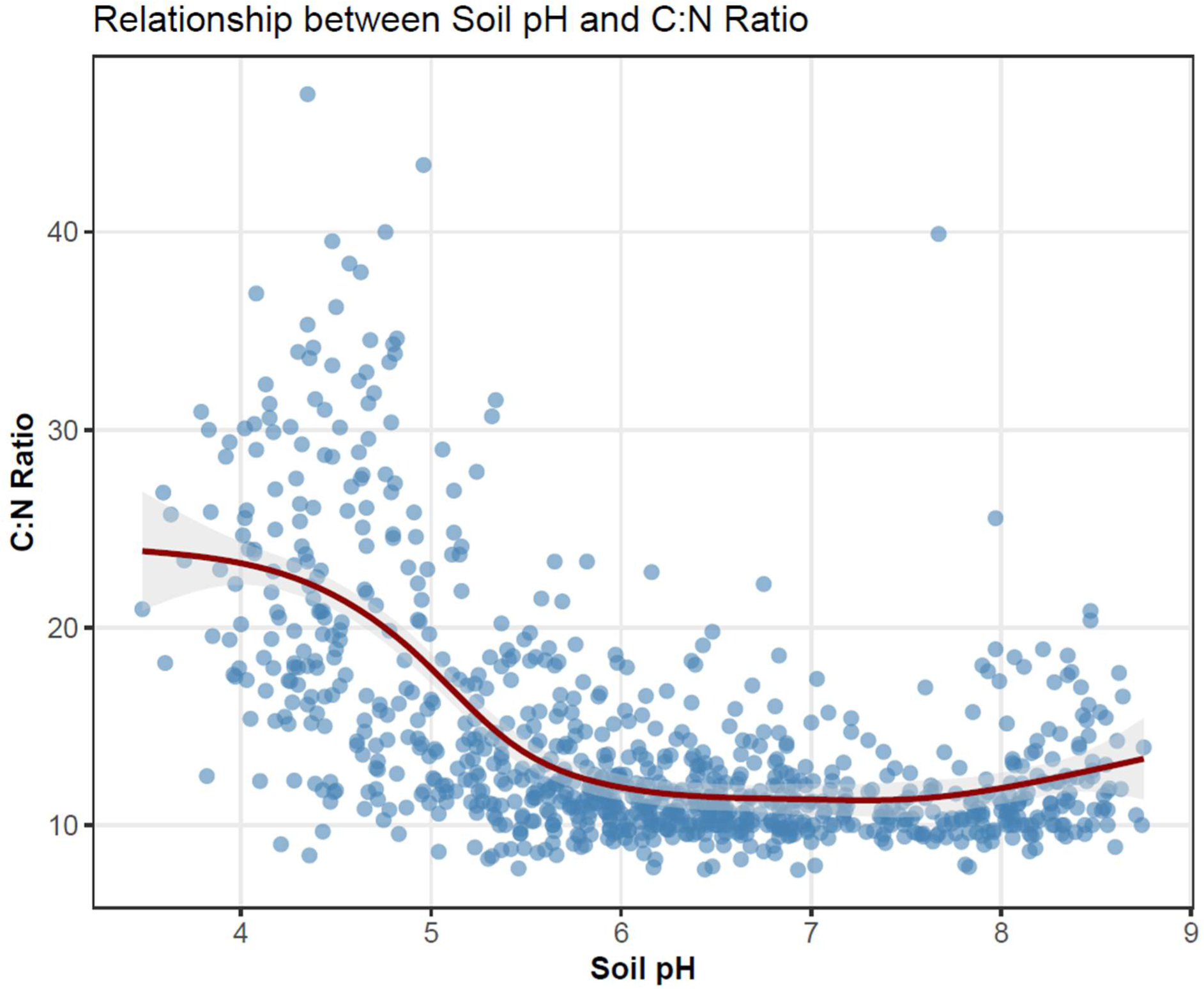
Relationship between the dominant hypervolume niche determinants

**Supplementary Table 1.**
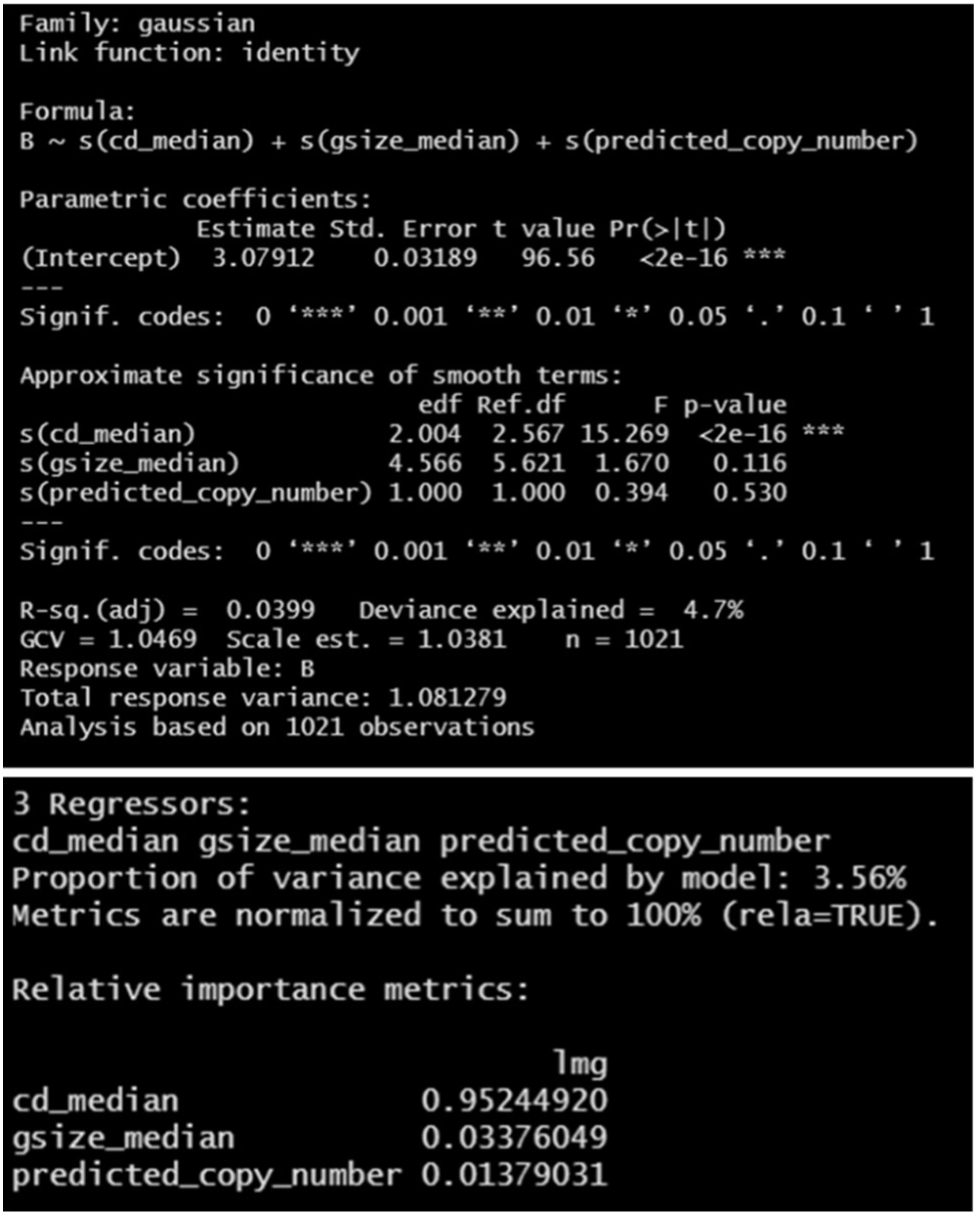
Gaussian Additive Model - library(mgcv) and Linear Model - library(relaimpo) findings relating genomic traits of ASV to Levins’ niche breadth score. cd_median = coding density; gsize_median = genome size; predicted_copy_number = rRNA copies; B = Levins’ niche breadth score.

**Supplementary Table 2.**
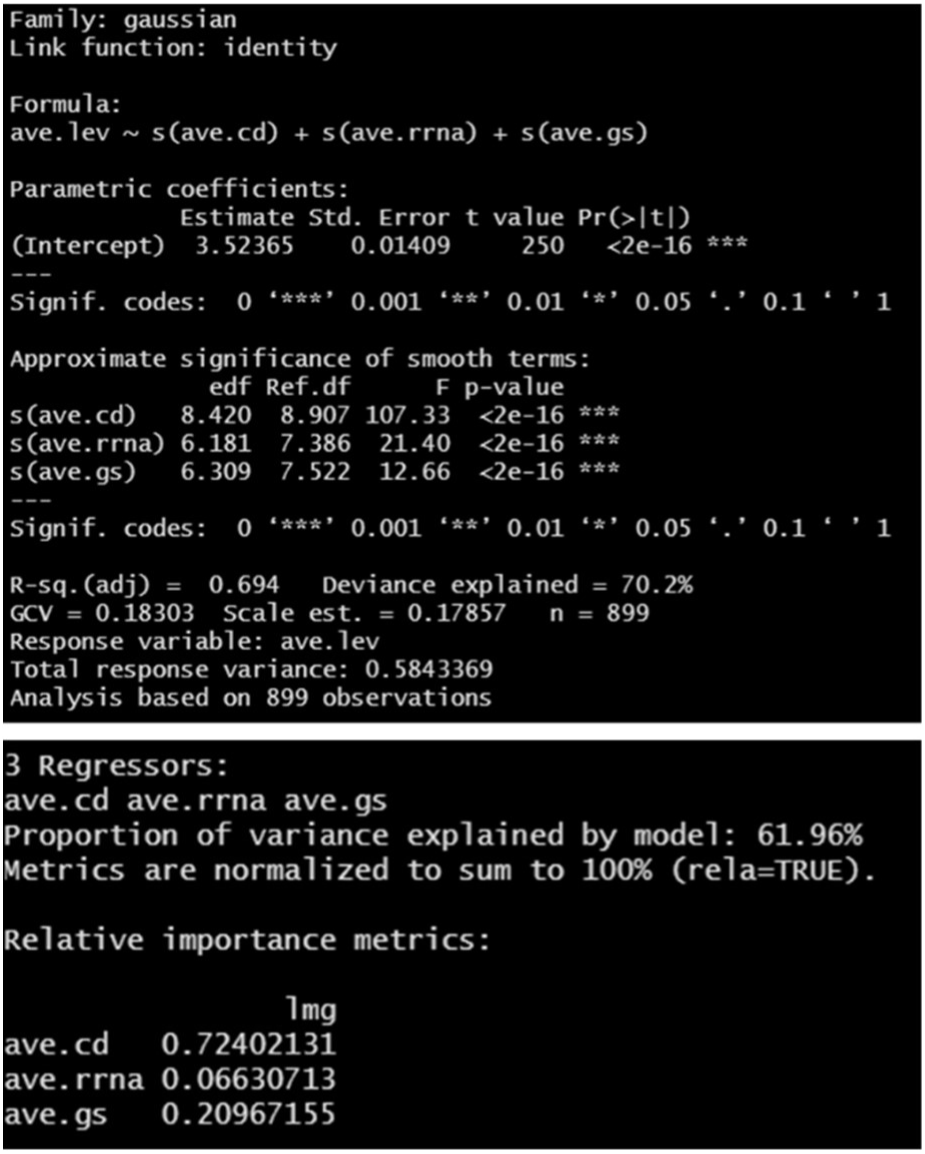
Gaussian Additive Model - library(mgcv) and Linear Model - library(relaimpo) findings relating community weighted mean (CWM) genomic traits to CWM Levins’ niche breadth score. ave.cd = CWM-Coding density; ave.rrna = CWM-rRNA copies; ave.gs = CWM-Genome size; ave.lev = CWM-Levins’ niche breadth score.

## Notes

### Competing Interest Statement

The authors have declared no competing interest.

### Funding

- UK Research and Innovation: BBX011089/1, NE/S005137/1

